# The antigenic landscape of N1 neuraminidase in human influenza A virus strains isolated between 2009 and 2020

**DOI:** 10.1101/2024.12.10.627724

**Authors:** João Paulo Portela Catani, Anouk Smet, Tine Ysenbaert, Daria Fijalkowska, Laura Amelinck, Yvonne Chan, Dan Tadmor, Philip Davidson, Satyajit Ray, Eric Camire, Liqun Han, Jianxin Zhang, Guadalupe Cortés, Katherine Roebke, Bianca Baum, John Hamberger, Maryann Giel-Moloney, Raul Gomila, Xavier Saelens, Thorsten U. Vogel

## Abstract

The immunogen composition of seasonal influenza vaccines is revised biannually to match the antigenic drift of the hemagglutinin of circulating influenza virus strains. Antibodies directed against the influenza neuraminidase also correlate with protection against influenza, yet the antigenic evolution of influenza neuraminidase remains underexplored. To evaluate the antigenic diversity of N1 neuraminidase, we generated a panel of immune sera in mice directed against 17 N1 neuraminidases derived from human H1N1 strains that were isolated between 2009 and 2020 and determined the neuraminidase inhibition (NAI) titers of these sera against a panel of 14 HxN1 viruses. The selected N1 neuraminidases cluster into two major phylogenetic groups while the NAI pattern revealed two antigenic groups corresponding to early and more recent H1N1 viruses. Sera raised against early N1 NAs inhibited NA activity from both early and recent H1N1 viruses equally well, whereas sera raised against more recent N1 NAs showed stronger NAI against recent H1N1 strains, indicating an asymmetric antigenic relationship between the two antigenic groups. Random forest modeling of antigenic distances indicated that the I321V, N386K/S and K432E substitutions played an important role in the N1 NA antigenic variation. Reverse-genetics experiments using mutant strains revealed that the N386K substitution in an early H1N1 virus increased susceptibility to inhibition by sera raised against recent N1 NAs, consistent with a role for the loss of the N-linked glycosylation motif and the acquisition of a lysine at this position near the lateral surface of the NA head. However, reciprocal substitutions in a recent N1 background did not restore the early antigenic phenotype, indicating a unidirectional effect. This finding suggests that anti-NA responses elicited against early pandemic H1N1 viruses remained effective against circulating N1 variants up to at least 2020.

## Introduction

Seasonal influenza is estimated to cause up to half a million deaths every year and is associated with a high economic and clinical burden^1–5^. Influenza vaccines reduce the disease caused by influenza virus infection although their effectiveness has varied from 10 to 60%^6,7^. Seasonal influenza vaccines are composed of representative strains of H1N1, H3N2 and one or two influenza B lineages. The recommended vaccine strains are constantly evaluated and, if needed, updated twice a year, aiming to ensure antigenic match with the predominant circulating strains. The antigenic evolution of human influenza viruses is typically determined in hemagglutination inhibition tests, which captures the antigenic drift of the viral hemagglutinin (HA)^8,9^. The neuraminidase (NA) is the second major influenza A and B virus surface antigen. Like HA, NA is subject to antigenic drift, indicating that immune responses against NA exert a selective pressure on influenza A and B viruses^10^. Furthermore, antibodies against NA correlate with protection against influenza^11–16^. These findings have contributed to the renewed interest in the development of influenza vaccines that contain a standardized amount of NA antigen to try to improve the efficacy and breadth of protection^17,18^.

We previously reported the antigenic evolution of N2 NAs derived from human H3N2 strains that were isolated between 2009 and 2017^19^. That work expanded the knowledge of N2 antigenic diversity and identified at least four N2 antigenic groups. Furthermore, we demonstrated that the antigenic assessment of recombinant N2 NA in mice provides similar results to those obtained in ferrets, which is considered the gold standard small animal model to study immunity and pathogenesis of human influenza viruses.

Since its introduction into the human population, the H1N1 2009 pandemic (H1N1pdm) virus has been subjected to immune selection pressure, which has resulted in antigenic drift of HA and NA^20^. Aiming to obtain a high-resolution insight into the antigenic relatedness of NAs derived from H1N1pdm viruses that were isolated between 2009 and 2020, we selected N1 NAs derived from 17 human H1N1 strains that were isolated between 2009 and 2020. The corresponding recombinant NAs were produced and used to immunize mice. NA inhibition (NAI) activity of the resulting sera was then determined against a panel of H1N1 and H6N1 viruses. NAI data analysis revealed that H1N1 viruses isolated between 2009 and 2020 could be classified into 2 antigenically distinct groups. A machine learning model was generated, which suggested an important impact of amino acid substitutions I321V, N386K and K432E on N1 NA antigenicity. The rescue of single mutant viruses revealed that removal of the N-glycosylation site by the substitution N386K was associated with unidirectional increased susceptibility to NAI by sera raised against NA from the more recent N1 phylogenic group.

## Results

### Immunogenicity of recombinant N1 NAs

Based on amino acid sequence diversity, and using minimax linkage clustering^21^ implemented in R protoclust package, 17 representative N1 NAs were selected from the GISAID database^22^ from human H1N1 viruses isolated between 2009 and 2020. The phylogenetic relatedness, based on the amino acid sequences of these selected N1 NAs, together with the NA of 5 other randomly selected H1N1 isolates per year from 2009 to 2020, is shown in Figure 1a. The selected N1 NAs belong to two major phylogenetic groups (referred to hereafter as PG1 and PG2) and are further divided in different clades confirming that minimax linkage clustering selected a diverse panel of N1 representatives. Amino acid differences in the head domains of the N1 NA panel are listed in supplementary table 1.

**Figure 1.**
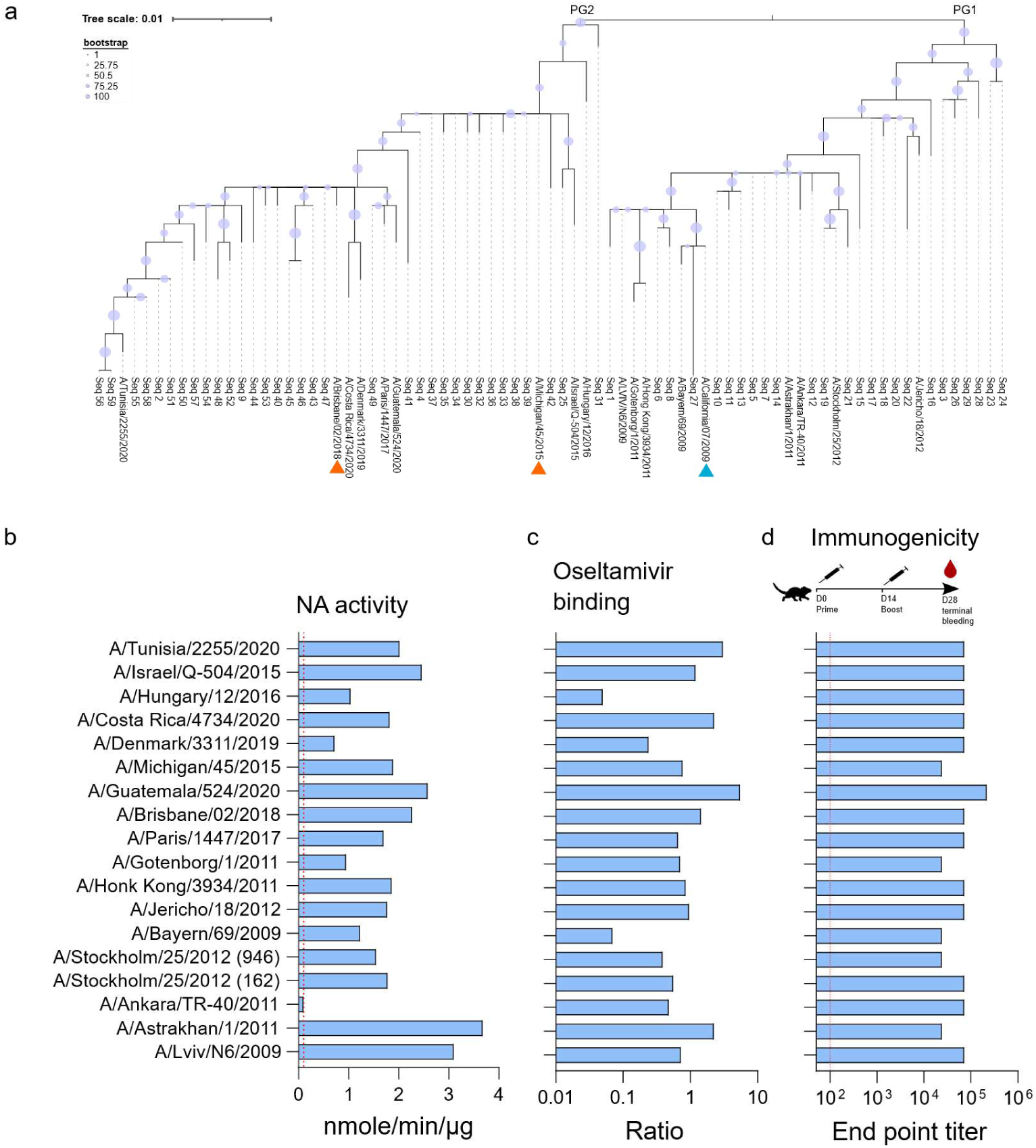
N1 NA selection, characterization of recombinant N1 NAs, and generation of immune sera. (a) Phylogenetic relatedness based on the amino acid sequence of 17 selected N1 NA head domains next to 5 randomly selected NA sequences from H1N1 viruses isolated between 2009-2020 (5 per year), and the A/California/07/2009 strain. Orange triangles: seasonal vaccine strains that were included here as N1 NA antigens; blue triangle: prototypic H1N1pdm. (b) The specific activity of the recombinant N1 NAs based on the MUNANA assay. (c) Ratio of protein to oseltamivir binding (mg/ml)/(mg/ml). (d) Groups of 5 female BALB/c mice were immunized in a prime-boost regimen with a 2-week interval with AF03-adjuvanted N1 NAs. Homologous endpoint ELISA titers were determined in pooled sera collected 4 weeks after the prime immunization.

Next, we generated recombinant N1 NA immunogens based on the 17 selected H1N1 viruses. These immunogens consisted of a tetramerizing tetrabrachion zipper fused at the N-terminus of the NA catalytic head and were expressed as secreted proteins by transfected CHO cells, as previously described.^19^ The specific activity of the purified N1 NA proteins was determined using 4-methylumbelliferyl-N-acetylneuraminic acid (MUNANA) as a substrate. All N1 NAs were active, except the NA derived from A/Ankara/TR-40/2011 (Figure 1b). All 17 N1 NAs could bind oseltamivir, indicating that the active site in the recombinant NAs was correctly folded (Figure 1c). Two batches (named 946 and 162) of N1 NA derived from A/Stockholm/25/2012 were produced and characterized, with comparable results (Figure 1b and 1c).

Next, groups of 5 mice were immunized with one µg of N1 NA, adjuvanted with AF03, in a prime-boost regimen with 2 weeks interval to generate immune sera. Two weeks after the last immunization mice were terminally bled, and sera were pooled per group. Analysis of the sera by ELISA using the respective recombinant N1 NAs as coated immunogens showed that all groups had seroconverted (Figure 1d).

### NA inhibition-based antigenic analysis reveals two distinct N1 NA antigenic groups

The NAI activity of each of the N1 NA mouse immune sera was determined using an enzyme-linked lectin assay (ELLA) against 14 different HxN1 viruses: H6N1 reassortants with N1 derived from A/California/07/2009, A/Michigan/45/2015, A/Hubei Wujiagang/SWL310/2013, A/Cameroon/9766/2017, A/Oman/5532/2017, or A/Wisconsin/588/2019; and H1N1 viruses A/Brisbane/02/2018, A/Hawaii/70/2019, A/Wisconsin/588/2019, A/Lviv/N6/2009, A/Stockholm/25/2012, A/Israel/Q-504/2015, A/Hungary/12/2016, and A/Ireland/84630/2018. The NAI titers were first normalized to the respective ELISA values and then relative to the IC_50_ value of the serum with the highest NAI titer for a given HxN1 virus and plotted as a heatmap, with columns indicating the influenza virus strains used as a source of NA in ELLA and rows representing the sera raised against the respective recombinant N1 NAs (Figure 2). The serum samples are ordered according to the phylogenetic relatedness of the recombinant N1 NAs that were used as an immunogen. Hierarchical clustering of the tested HxN1 strains revealed two major N1 NA antigenic groups: a first group of strains from 2009-2013 (AG1) and a second group comprising strains from 2015-2020 (AG2). The antigenic clustering accords very well with the two phylogenetic groups. Plotting of the raw NAI data (i.e. NAI titers that were not normalized based on maximal IC_50_ values), showed that immune sera raised against N1 NAs belonging to PG1 had similar NAI titers against viruses from AG1 and AG2. Conversely, sera raised against N1 NAs classified in PG2 displayed stronger inhibition of strains from AG2 (Figure 2 – Figure supplement 1).

**Figure 2.**
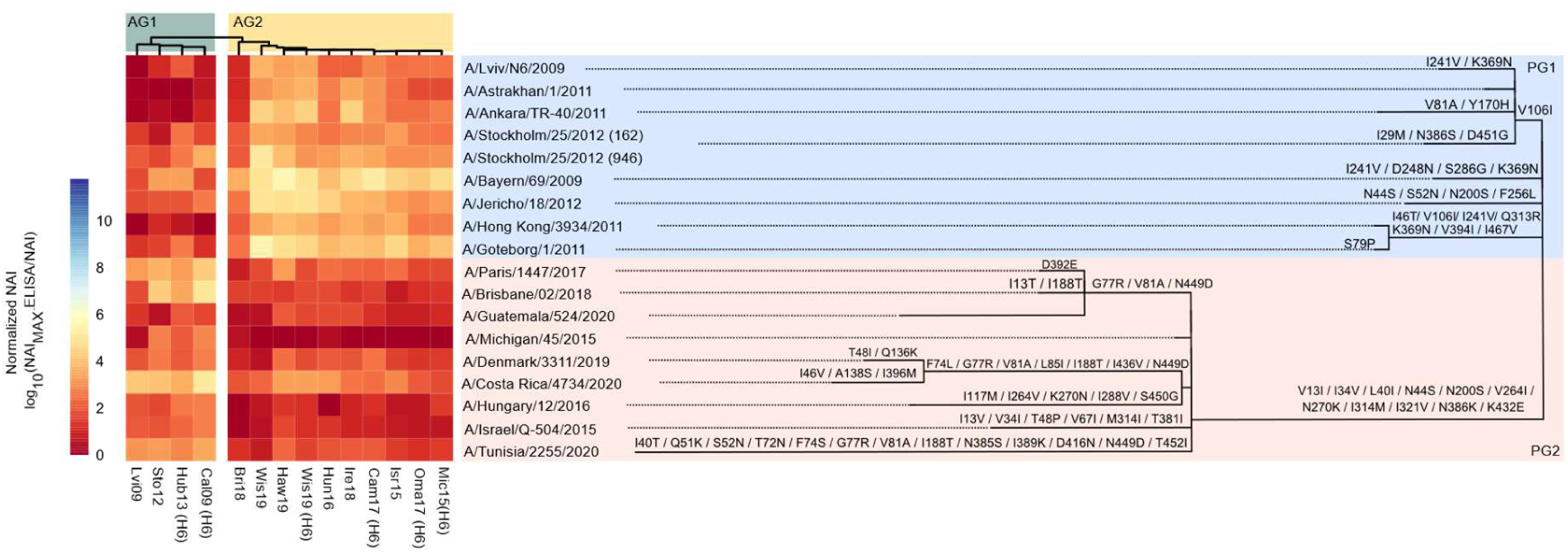
Breadth of N1 NA inhibition. NAI against 14 different HxN1 strains (columns) by mouse sera raised by immunization with recombinant N1 NA (rows). The sera are ordered according to the phylogenetic relatedness of the N1 NA head domain amino acid sequences, with amino acid substitutions depicted above the branches. The two phylogenetic groups are color coded and labeled as PG1 and PG2. Hierarchical clustering is depicted above the heat-map with the derived antigenic groups AG1 and AG2 indicated. The red to blue scale represents the normalized NAI values for each virus as fold lower than the serum with the highest normalized NAI for each virus.

Comparison of the normalized NAI IC_50_ titers of the sera raised against the 2 batches of N1 A/Stockholm/25/2012 (162 and 946) revealed a significantly lower NAI of batch 946 immune serum against all but one of the HxN1 viruses (Figure 2 – figure supplement 2a). Despite this difference the antigenic profile of the sera raised against the two A/Stockholm/25/2012 N1 NA batches is highly similar (R^2^=0.86) (Figure 2 – figure supplement 2b).

Visualization of the antigenic differences using multidimensional scaling according to Smith *et al*.^23^, showed a spatial distribution that is in line with the hierarchical clustering (Figure 2 – figure supplement 3).

### A Machine Learning tool to predict N1 NA antigenic distances and relevant amino acid substitutions

Next, we calculated the sera-to-sera average antigenic differences and used these differences in a random forest (RF) modeling (Figure 3a). Similar to Yao et al.^24^, pairs of amino acid substitutions were featured as index values from scoring matrices or, in addition, overall sequence similarity. The mtry parameter (i.e. the number of tested conditions per node) of the RF model was optimized for each amino acid index matrix (Figure 3 – figure supplement 1). RF was trained using a 5-fold cross-validation data set, repeated 10 times. The performance of each matrix was measured as R^2^ from predicted versus observed antigenic differences. The highest correlation coefficient was obtained from the matrix HENS920103, with an R^2^ of 0.762 (Figure 3b). When compared to all other tested matrices this outcome did not differ from the ATLS910101, BENS940102, BENS940101 and HENS920102 matrices (Figure 3 – figure supplement 2). Figure 3 – figure supplement 3, illustrates a single decision tree extracted from the RF model. The most interesting aspect of RF is the ability to score the importance of particular features. The increase in mean square error (MSE) is an importance score that can be calculated by randomly permuting values of a feature (here amino acid substitutions or overall similarity). Figure 3c represents the weighted average of importance scores from the top 6 best performing matrices, and highlights that substitutions at positions 321, 432 and 386 are most likely to impact NAI. Interestingly, the amino acid residues at these positions are surface exposed with amino acid 432 positioned close to the catalytic pocket of N1 NA, while amino acids 321 and 386 are located at the lateral side of N1 NA (Figure 3d).

**Figure 3.**
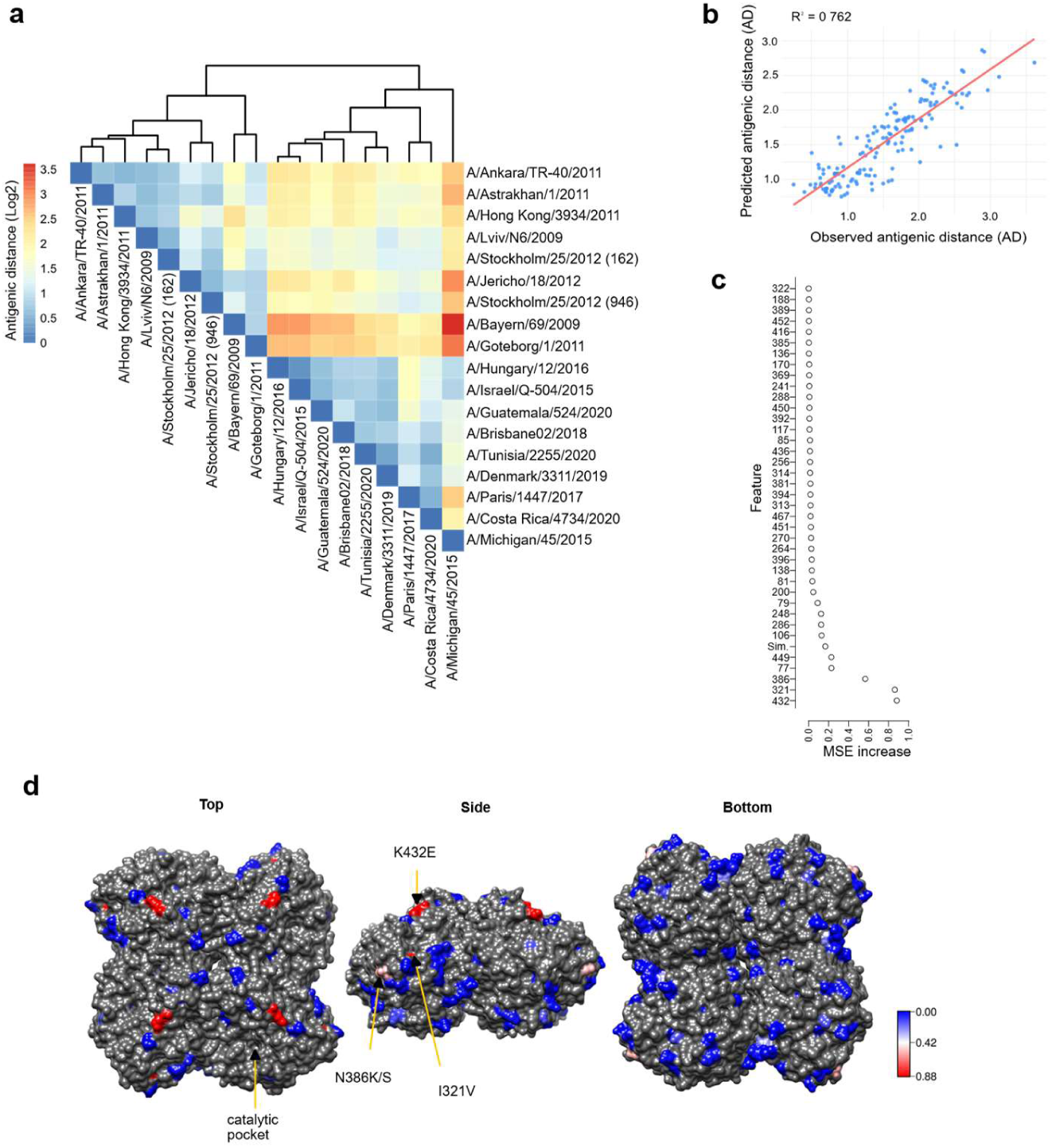
Machine learning algorithm allows to predict the antigenic distance and antigenically important amino acid substitutions. (a) Sera to sera differences were used to calculate the average antigenic distances, which are represented in the heat map. (b) Correlation of the antigenic distances predicted by the RF model and the observed values. (c) The Mean Squared Error Increase (MSEI) for each amino acid substitution in the N1 NAs obtained with the best performing amino acid index matrices. (d) Distribution of amino acid substitutions on the surface of N1 NA. The substitutions are indicated in red to blue scale representing MSEI values. Top, side, and bottom views of tetrameric H1N1pdm 2009 NA (PDB: 3NSS) are shown. The NA catalytic pocket is indicated on one of the protomers on the top view of the structure.

We next evaluated the impact of amino acid substitutions at positions 321, 386, and 432 on susceptibility to NAI by the N1 NA immune serum panel. Site directed mutagenesis was used to introduce I321V, N386K and K432E in the NA of A/Belgium/ma-145/2009 (Bel09), a strain belonging to AG1. In addition, the substitutions V321I, K386N, K386S, V321I/K386N, and V321I/E432K were introduced in the NA of A/Wisconsin/588/2019 (Wis19), which belongs to AG2. H6N1 reassortant viruses carrying NA with these substitutions were successfully rescued and the NAI was determined using the N1 NA immune serum panel (Figure 4 – a and b). This analysis revealed that all mutant viruses, except N386K in Bel09 NA, had an antigenic profile similar to the parental antigenic group (Figure 4c). The H6N1 virus with Bel09-N386K NA was more susceptible to NAI by sera raised against PG2 NAs than H6N1 virus with parental Bel09 (Figure 4 – figure supplement 1).

**Figure 4.**
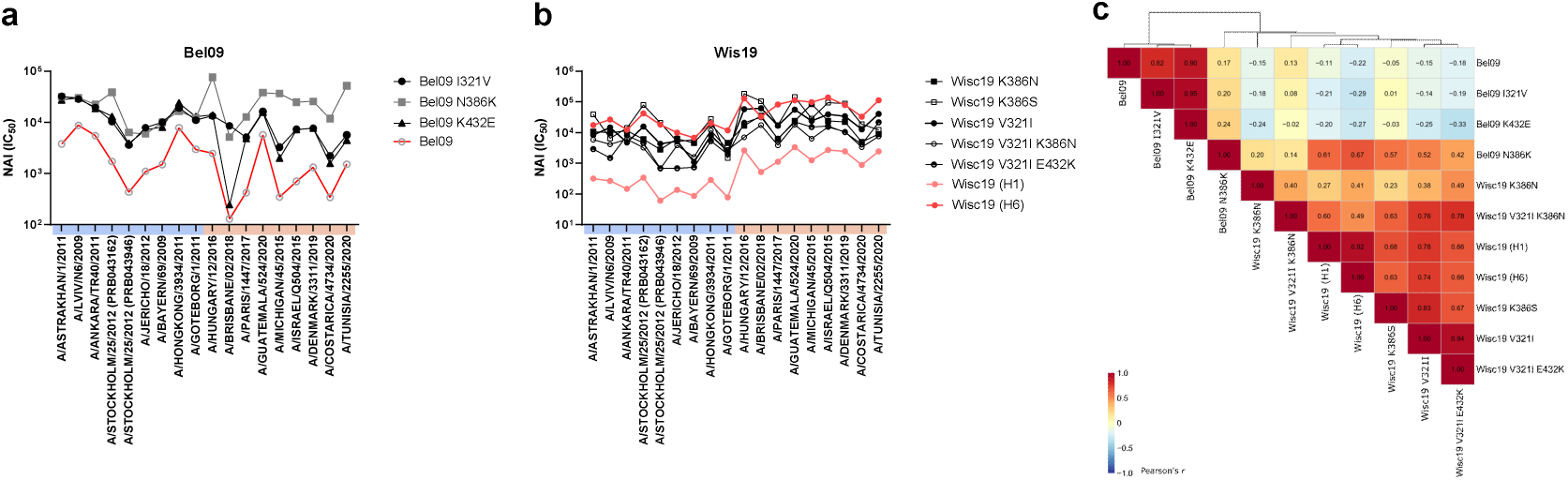
NAI determined against H6N1 viruses with substitutions in NA. (a) H6N_Bel09_ carrying WT N1 or N1 with substitutions I321V, N386K, or K432E. (b) HxN_Wis19_ carrying K386N, K386S, V321I, V321I/K386N, or V321I/E432K substitutions in N1, or the parental N_Wis19_ NA (as H6 or H1references, as shown in Figure 2). (c) Pearson correlation coefficients of NAI among mutants and parental strains.

## Discussion

The protective effect of anti-NA antibody responses against influenza has been known for many decades and repeatedly observed in clinical studies^14–16,25–27^. Nonetheless, currently used virus-derived influenza vaccines induce a poor response against NA^28^. Therefore, the addition of NA to influenza vaccines to improve their effectiveness has been proposed^29^. However, among other reasons, the labile nature of the NA tetramer and limited understanding of NA antigenic drift hamper major efforts to bring this concept to clinical reality^30^. Previously, using a panel of mouse and ferret sera and recombinant, soluble tetrameric NA immunogens we determined the antigenic landscape of N2 NA of human H3N2 influenza viruses isolated between 2009 and 2017^19^. Here we performed a similar analysis of human H1N1 NAs using immune sera from mice. We selected a genetically divergent panel of N1 NAs derived from human H1N1 viruses that circulated from 2009 until 2020. These N1 NAs were produced as soluble recombinant proteins, stabilized as a tetramer by a heterologous tetrabrachion domain. This strategy allows the induction of an immune response which is free of anti-HA antibodies and thus enabled us to capture the NA inhibition without potential interference by such antibodies. The characterization of the recombinant N1 NAs included the determination of enzymatic activity and binding to oseltamivir. The A/Ankara/TR-40/2011-derived recombinant N1 NA was the only inactive protein in the MUNANA assay, although it showed binding to oseltamivir and induced NAI antibodies, which indicates that, despite non-detectable enzymatic activity, the protein was correctly folded.

The inclusion of 2 independent groups of mice that were immunized with different preparations (batches 946 and 162) of NA from A/Stockholm/25/2012 revealed a degree of batch-to-batch variability. Batch 946 had lower enzymatic activity, oseltamivir binding ratio and induced slightly lower homologous ELISA titers compared to batch 162. Even after normalization of the NAI IC_50_ with the ELISA values, values were on average 0.66-fold lower in the immune sera raised against batch 946 compared to batch 162 of A/Stockholm/25/2012 N1 NA. This result points out that minor antigenic differences observed in different antigens may be masked by experimental biases.

NAI determination using the panel of 18 anti-N1 NA sera against 14 HxN1 viruses revealed that pandemic 2009-derived N1 NAs can be classified in at least 2 distinct groups. The AG1 comprises H1N1 viruses that circulated in the early years after the start of the 2009 H1N1 pandemic and in our panel is represented by strains from 2009 until 2013. The AG2 is formed by more recent strains, isolated between 2015 and 2020. The antigenic classification agrees well with the phylogenetic relatedness of the strains which is characterized by the substitutions V13I, I34V, L40I, and N44S in the NA stalk and N200S, V264I, N270K, I314M, I321V, N386K, and K432E in the catalytic head domain (Figure 2). All vaccine recommended strains since 2015 bear the full set of substitutions that characterize the antigenic group 2 (V13I, I34V, L40I, N44S, N200S, V264I, N270K, I314M, I321V, N386K and K432E). Antigenic distances based on sera-to-sera average differences was used to build a RF model that can predict the antigenic distance between two N1 NAs based on their amino acid sequences. This RF model also allows to predict which amino acid substitutions are most likely responsible for the antigenic differences. The substitutions V13I, I34V, L40I, and N44S are located in the N1 stalk, and thus absent in the recombinant tetrabrachion stabilized NAs. Therefore, these substitutions were not included in the RF analysis. Among the remaining substitutions, based on the RF model, I321V, N386K/S, and K432E stand out as the most important ones (Figure 3c). In agreement with our previous study on N2 NA^19^, those substitutions are located close to the catalytic site and at the lateral face of NA.

Interestingly, I321V and K432E co-occur and are the only substitutions among the two antigenic groups of our tested N1 NAs that did not revert to the ancestral residues I321 and K432. To try to untangle the contribution of those substitutions we rescued the H6N1 viruses with the NA of A/Belgium/ma-145/2009 (Bel09) bearing the substitutions I321V, N386K, or K432E and determined their susceptibility to the N1 NA immune serum panel. The NAI values indicate that the N386K substitution, which removes a N-glycosylation site, slightly increases the recognition of NA by sera raised against NAs of PG2, while maintaining equal susceptibility to AG1 immune sera. Gao *et al*. reported that the loss of an N-linked glycosylation site at position 386 in NA of H1N1 viruses that circulated in the 2017-2020 seasons contributed to an antigenic change in NA compared to NA from H1N1 viruses that circulated in the early years after the H1N1pdm 2009 pandemic outbreak ^31^. This conclusion was based on NAI titers of ferret sera raised against H1N1 strains from 2015, 2018, and 2019 against PR8 single gene reassortant viruses carrying the respective CA09 NA mutants. The presence of an N-glycosylation site at position 386, reduced recognition of CA09 N1 by sera raised against more recent N1 NAs and not the other way around, which is in line with our observations (Figure 2 – Figure supplement 1).

Using a panel of seven monoclonal antibodies isolated from individuals that had been infected with H1N1pdm, Yasuhara *et al*., reported the antigenic distinction between the NA of the prototype A/California/04/09 strain (clade 1) and more recent isolates such as A/Yokohama/94/2015 (clade 6B.1). The substitutions surrounding the catalytic pocked, including the substitution K432E, were described to impact binding of monoclonal antibodies and reduced NAI in some of the human sera collected in 2009-2010 ^32^. In our hands the substitution of K436 alone was not sufficient to alter the NAI profile of a panel of polyclonal mouse immune sera, which may indicate that multiple substitutions are required to promote N1 evasion from a polyclonal humoral response. Furthermore, our study was performed with immune sera raised in naïve mice whereas the exposure history of the influenza NA antigen in the human population is much more complex, which may influence the quality and intensity of the elicited NAI, and explain some of these discrepancies.

Interestingly, the N386S mutation occurs in A/Stockholm/25/2012, which also results in loss of the N-glycosylation site, yet did not result in higher NAI titers against recent N1s, suggesting that the loss of an N-glycosylation site at position 386 alone does not explain the major antigenic difference between early G1 and the recent G2 H1N1 viruses. We hypothesize that the presence of a positively charged lysine residue at position 386 (and not the polar serine residue) may favor epitope recognition by B-cells, a hypothesis which remains to be further investigated.

We have also rescued the AG2 Wis19 strain bearing the K386N, K386S, V321I, V321I/K386N, or V321I/E432K substitutions. None of those substitutions, however, reversed the NAI profile of Wis19 to AG1. This result confirms the unidirectional impact of substitution at position 386, which suggests no major relevance in the antigenic drift.

Although our study focused on NAI, antibodies that bind NA in absence of measurable inhibitory activity can also contribute substantially to protection via Fc-mediated effector mechanisms^33^, indicating a broadened protective potential of anti-NA antibody response^34^. While the majority of clinical studies have reported NAI as a correlate of protection^12–16,26,27,35^, a few have investigated NA-biding antibodies ^36^.

In conclusion, by analyzing NAI of a diverse panel of anti-H1N1 NA immune sera against HxN1 NA viruses, our study identified that serum raised against N1 NAs from PG1 exhibited similar NAI titers against earlier AG1 and recent AG2 strains. Immune sera raised against recent PG2 N1 NAs, however, had higher NAI titers against AG2 than against AG1 strains. This unidirectional behavior is majorly characterized by increased NAI titers induced by PG2 N1 NAs against AG2, and thus, does not reflect escape from AG1 NA immunity in the population. This observation is aligned with the lost N-glycosylation site and may be exacerbated by increased epitope recognition due to presence of lysine in proximity of the catalytic pocked. This result suggests that the anti-NA response against early pandemic H1N1 viruses would remain effective against H1N1 viruses that circulated up until 2020.

## Materials and methods

### Design and production of recombinant proteins

The CD5 secretion signal, an amino-terminal His-tag and the thrombin cleavage site followed by tetrabrachion domain, were fused to the N1 NA head domain (aa 74-469) coding sequence and cloned under the transcriptional control of the CMV promoter in the pCDNA3.4 plasmid. The recombinant tetNAs were expressed in a mammalian cell culture system as previously described with modifications^19^.

### Virus generation and amplification

The HxN1 read out panel viruses are 6:2 reassortants on an A/Puerto Rico/8/1934 backbone (PB2, PB1, PA, NP, M, NS). The NA gene from the desired strain was matched with an HA gene matching the H1N1 strain or derived from H6 A/Mallard/Sweden/81/2002. The eight influenza segments were cloned in the bidirectional plasmid pHW2000S, and the resulting plasmids were transfected into HEK293FT cells using a lipid-based transfection reagent^37^. HEK293FT cells were co-cultured with MDCK ATL cells during transfection, or MDCK ATL cells were added 24hrs post transfection of HEK293FT cells to allow for efficient amplification of Influenza virus. The virus harvest was done 4 to 6 days post transfection, (this initial recovery stage referred to as P0). Harvested P0 virus was further passed on MDCK ATL cells to generate P1 and P2 working virus stocks that were characterized for infectious titer and NA sequence confirmation. Working virus stocks (P2) were passaged on specific pathogen free (SPF) white leghorn chicken eggs to further amplify virus stocks. Harvested allantoic fluid was clarified by centrifugation and characterized for infectious titer, NA gene sequence identity and NA enzymatic activity.

### NA enzymatic activity

The enzymatic activity of recombinant tetNAs was determined with the fluorogenic small molecule substrate 4-methylumbelliferyl-N-acetylneuraminic acid (MUNANA) as follows. Ten microliters of 1 mM of 4-MUNANA (Sigma cat # M8639) in 200 mM sodium acetate buffer (pH 6.5) containing 2 mM CaCl_2_ and 1% butanol was incubated with 40 µl of PBS solution containing 30, 15, 7.5, or 3.75 ng of tetNA. Conversion to 4-methylumbelliferone (4-MU) was monitored every 2 min for 1 h and quantified with an Infinite 200 Pro Tecan reader (Tecan Life Science) (excitation at 365 nm and emission monitored at 450 nm). A standard curve with 800 to 25 pmoles/well of 4-MU (Sigma cat # M1508) was used to extrapolate the molar conversion of MUNANA per microgram of purified recombinant tetNAs.

### Oseltamivir-NA binding assay using Octet Bio-Layer Interferometry

Binding of NA to Oseltamivir Phosphate-Biotin (Tamiflu-Biotin, GlycoTech Inc.) was measured on the Octet-HTX (ForteBio, Sartorius). All measurements were performed in Sartorius Kinetic Buffer (1xKB buffer containing PBS, 0.02% Tween20, 0.1% BSA, and 0.05% sodium azide). The Tamiflu-Biotin was immobilized on streptavidin biosensors (ForteBio, Sartorius) at a concentration of 0.25-0.5 µg/ml in 1xKB buffer. Binding of NA to Oseltamivir was initiated by dipping Oseltamivir bound biosensors into sample wells containing a 2-fold dilution series of recombinant NA (0.8–200 nM in 1xKB buffer) at 25 °C with a 5 min association step and a 5 min disassociation step. The level of binding response is proportional to the concentration of recombinant NA. The response at the end of the association step was used for determination of kinetics, affinity, and quantitation regarding the NA bound to the biosensor tip.

### Mouse experiments

The mouse immunization experiments were conducted according to the Belgian legislation (Belgian Law 14/08/1986 and Belgium Royal Decree 06/04/2010) and European legislation on protection of animals used for scientific purposes (EU directives 2010/63/EU and 86/609/EEC). Experimental protocols were approved by the Ethics Committee of the Vlaams Instituut voor Biotechnologie (VIB), Ghent University, Faculty of Science (EC2022-056). Female BALB/c mice, aged 6–7 weeks, were purchased from Janvier. The mice were housed in a specified pathogen-free animal house with food and water *ad libitum*. Animals, under isoflurane anesthesia, were immunized twice, with an interval of 14 days, by intramuscular injection in the right quadriceps. Immunizations containing 1 µg of recombinant protein were performed in a volume of 50 µl (25 μl of antigen in PBS + 25 μl of AF03 per dose)^38^. Two weeks after the boost, mice were humanely sacrificed by an overdose of pentobarbital, delivered intraperitoneally (6mg/mouse), and bled retro-orbitally. The obtained blood samples were incubated for 30 min at 37°C to allow clotting, which was followed by centrifugation at 10,500×*g* for 5 min. The supernatant (the serum) was recovered and submitted to a second centrifugation at 10,500×*g* for 5 min. Cleared sera were stored at –20°C before use in serological assays.

### ELISA

Anti-tetNA IgG titers in mouse sera were determined by capture ELISA using tetNAs in Pierce nickel-coated plates (cat. # 15442). Recombinant proteins were diluted to 0.5 µg/ml in DPBS (Life Technologies cat. # 14040-182). Then, 50 µl of the coating antigen solution was added to each well and the plate was incubated at room temperature for 1 h on a shaking platform. The plates were then washed three times with PBS-T (Sigma cat. # P3563-10PAK) and blocked for 1 h with 1% BSA. After blocking, the plates were washed once with PBS-T and incubated with a threefold serial dilution, starting at a 1/100 dilution, of serum in DPBS, 0.5% BSA, 0.05% Tween-20 for 2 h at room temperature on a shaking platform. The plates were then washed five times with PBS-T and incubated with a 1:5000 dilution of anti-mouse IgG-HRP (GE Healthcare cat. # NA931-1ml) in DPBS, 0.5% BSA, 0.05% Tween-20. The 3,3’,5,5’-tetramethylbenzidine (TMB) substrate (BD cat. # 555214) was added after three washes with PBS-T and the reaction was stopped after 5 min by addition of 50 µl of 1 M H_2_SO_4_. The optical density (OD) in each well was determined at 450 nm and, as a reference, 655 nm using an iMark Microplate Absorbance Reader (Bio-Rad). The end point titer was determined for each serum sample by scoring the dilution that resulted in an OD that was equal to or two times higher than the background OD obtained from pre-immune control sera dilution series.

### ELLA to determine NAI titers

Fetuin (Sigma cat. # F3385) was diluted into coating buffer (KPL cat. # 50-84-01) to a concentration of 25 µg/ml and 50 µl was added to the wells of Nunc MaxiSorp plates (Thermo Fisher cat. # 44-2404-21), which were subsequently incubated overnight at 4°C. The coated plates were then washed three times with PBS-T (Sigma cat. # P3563-10PAK) and incubated overnight with 60 μl of a dilution of H6N1 or H1N1 and 60 µl of twofold serial dilution of serum, starting at a 1/20 dilution, in sample buffer (1× MES (VWR cat. # AAJ61979-AP): 20 mM CaCl_2_, 1% BSA, 0.5% Tween-20). These dilutions of the H6N1 and H1N1 viruses correspond to the 70% maximum activity of NA from the respective viruses as determined in the ELLA. Fetuin-coated plates were then washed three times with PBS-T and incubated for 1 h with a solution of PNA-HRP (cat. # L6135-1MG, Sigma) at 5 μg/ml in conjugate diluent (MES pH 6.5, 20 mM CaCl_2_, 1% BSA). The plates were washed three times with PBS-T, TMB substrate was added, and then the plates were incubated for 5 min before the reaction was stopped by the addition of 50 µl of 1 M H_2_SO_4_. The optical density was measured at 450 nm and, as a reference, 655 nm in an iMark Microplate Absorbance Reader (Bio-Rad). Half maximum inhibitory concentrations (IC_50_) values were determined by non-linear regression analysis (GraphPad Prism software). NAI values were expressed as normalized by ELISA AUC titers.

### Random Forest algorithm

The RF algorithm, based on Yao *et al*., 2017, was described previously with two key adaptations: the inclusion of amino acid index matrices and overall sequence similairy^19,24^. Briefly, the non-conserved positions in the multiple sequence alignment of N1 head domain amino acid sequences of all tested N1s were selected.

For each substitution matrix *M*(e.g., BLOSUM62, PAM250, Binary, and optionally AA index-derived matrices), we build a pairwise feature matrix *X*^(M)^ by scoring every strain pair (*i*, *j*) at each variable position *k*. We also compute an auxiliary Similarity feature per pair as the mean of the valid position scores for that pair. Let *m* be the number of variable positions retained from the alignment. Then each row of *X*^(M)^contains up to *m* position features plus the Similarity feature for a single strain pair: 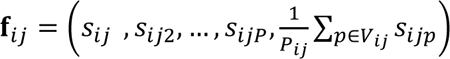, where f = matrix row, S = similarity at position *m, V* the set of variable positions and *Pij* the number of positions.

The response vector (Y) is composed of the antigenic distances for each pair of tested sera, computed as the log2 drop from the maximum titer across sera, then taking the average absolute difference between two antigens across all sera. Thus *Y*_ij_ reflects the serum-averaged antigenic separation of strains *i* and 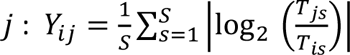, where *T*_is_ is the titer of antigen *i* against serum *s,* and *S* the number of sera pairs.^39^

For each matrix *M*, we fit a Random Forest regressor^40^ (package randomForest via caret) using repeated K-fold cross-validation. We fix *K* = 5 folds and 10 repeats, precompute and reuse the same resampling indices for all matrices. The only tuned hyperparameter is mtry (number of candidate predictors considered at each split). For each resample we save the out-of-fold predictions and then pool all held-out predictions to compute cross-validated metrics per matrix: *R*^2^as cor(ŷ, *y*)^2^and RMSE as 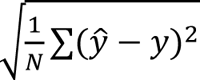. Feature importance at the RF level is summarized by the permutation-based %IncMSE returned by r Random Forest. The R script was prepared with assistance of Microsoft Copilot (Mar 30, 2026); all content was revised by a bioinformatician and can be found at https://github.com/SaelensLAB/RF_N1_eLife.

### Antigenic cartography

Antigenic cartography was performed as described by Smith *et al*., 2004, using the R CRAN package (https://CRAN.R-project.org/package=Racmacs)^23^. The antigenic map was optimized 5000 times, and the best optimization was kept (lowest stress value). Uncertainty was measured by bootstrapping 1000 repeats with 100 optimizations per repeat.

## Author Contributions

JPPC, TY, AS, LA, MGM, JH, BB, KR, GC, JZ, LH, EC, SR, PD, DT, YC performed experimental work. JPPC, TUV, and XS designed the experiments, analyzed and interpreted the data. DF revised the RF modeling. JPPC and XS wrote the manuscript and are accountable for accuracy and integrity of the work. All authors proofread and critically reviewed the manuscript.

---

**Figure 2 - Figure supplement 1.**
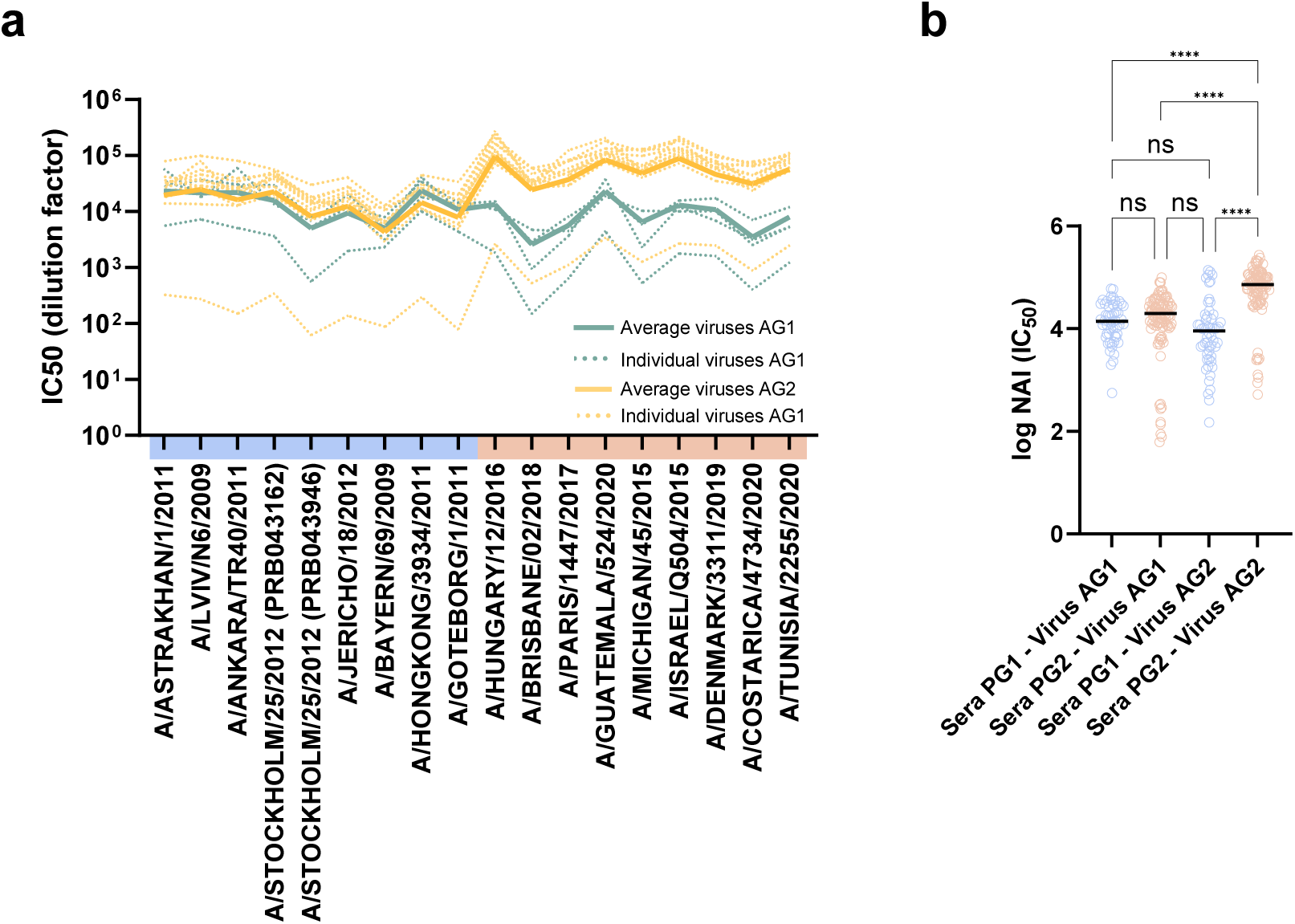
NAI across distinct antigenic groups. (a) Raw (non-max normalized) NAI titers (Y-axis) of serum samples, colored according to their phylogenetic relatedness (X-axis). Dotted lines represent NAI titers against individual viruses used as sources of NA, colored by antigenic group (AG1, blue; AG2, yellow). Solid lines indicate the average NAI values across viruses within the same antigenic group. (b) Comparison of NAI values grouped by serum and virus antigenic groups.

**Figure 2 – Figure supplement 2.**
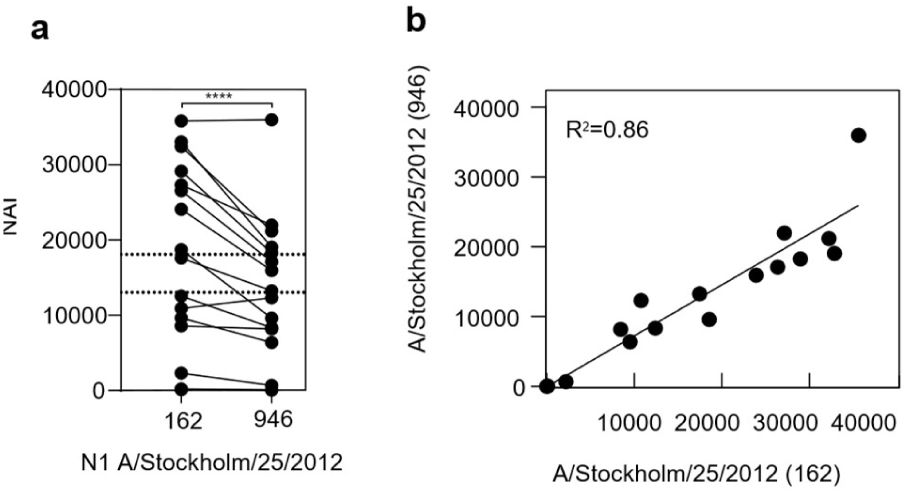
Comparison of immune serum raised against two batches of recombinant A/Stockholm/25/2012 N1 NA. (a) Paired dot plot showing IC_50_ values and (b) scatter plot with linear regression reporting Pearson correlation of ELISA normalized NAI of sera raised against batch 162 and 946 of recombinant N1 NA derived from A/Stockholm/25/2012. ****P<0.0001, determined with a paired t test.

**Figure 2 – Figure supplement 3.**
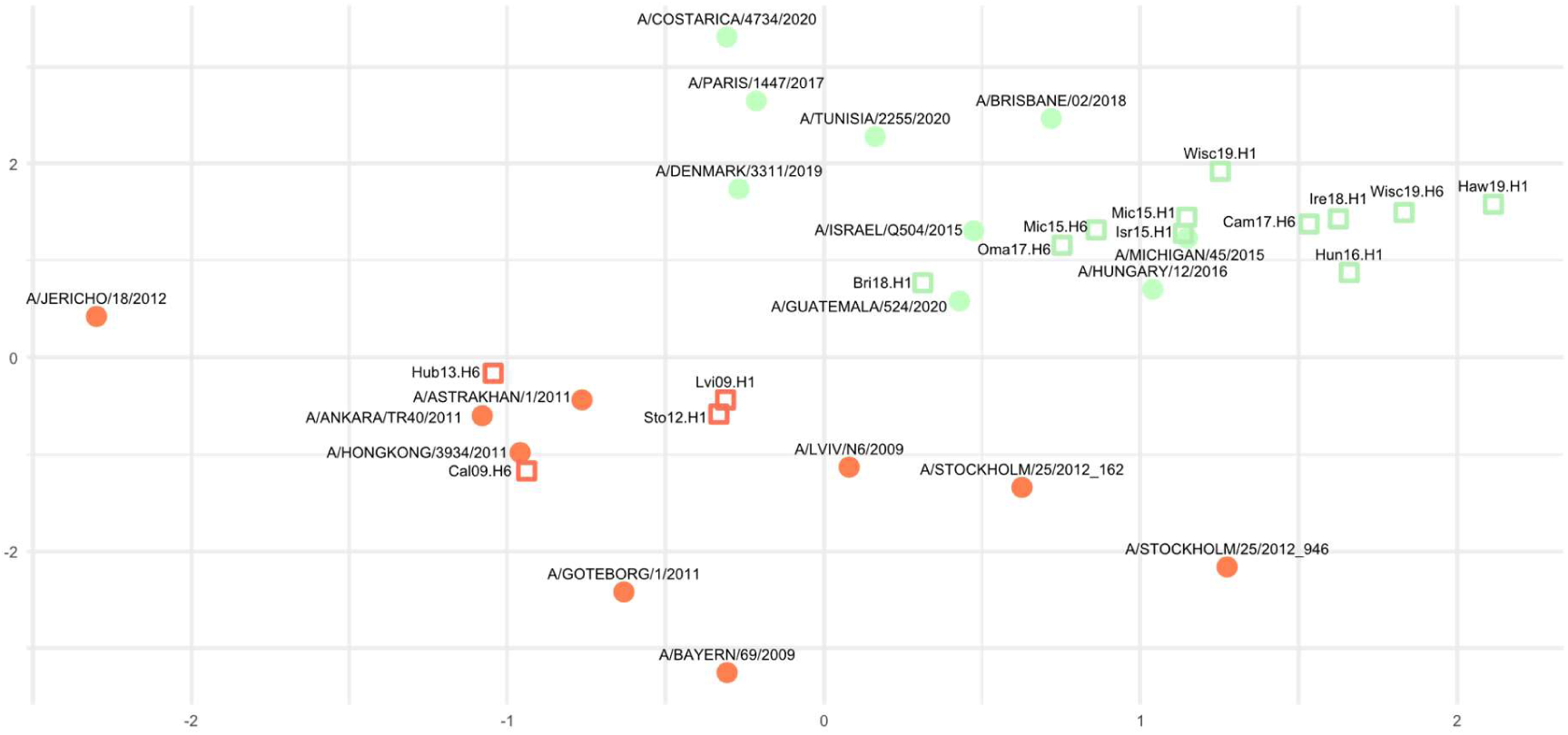
Antigenic map of NA of the H1N1 viruses. Sera are represented by dots and virus strains by squares. The phylogenetic and antigenic group 1 NAs are represented by orange and phylogenetic and antigenic group 2 NAs by green symbols.

**Figure 3. Figure supplement 1.**
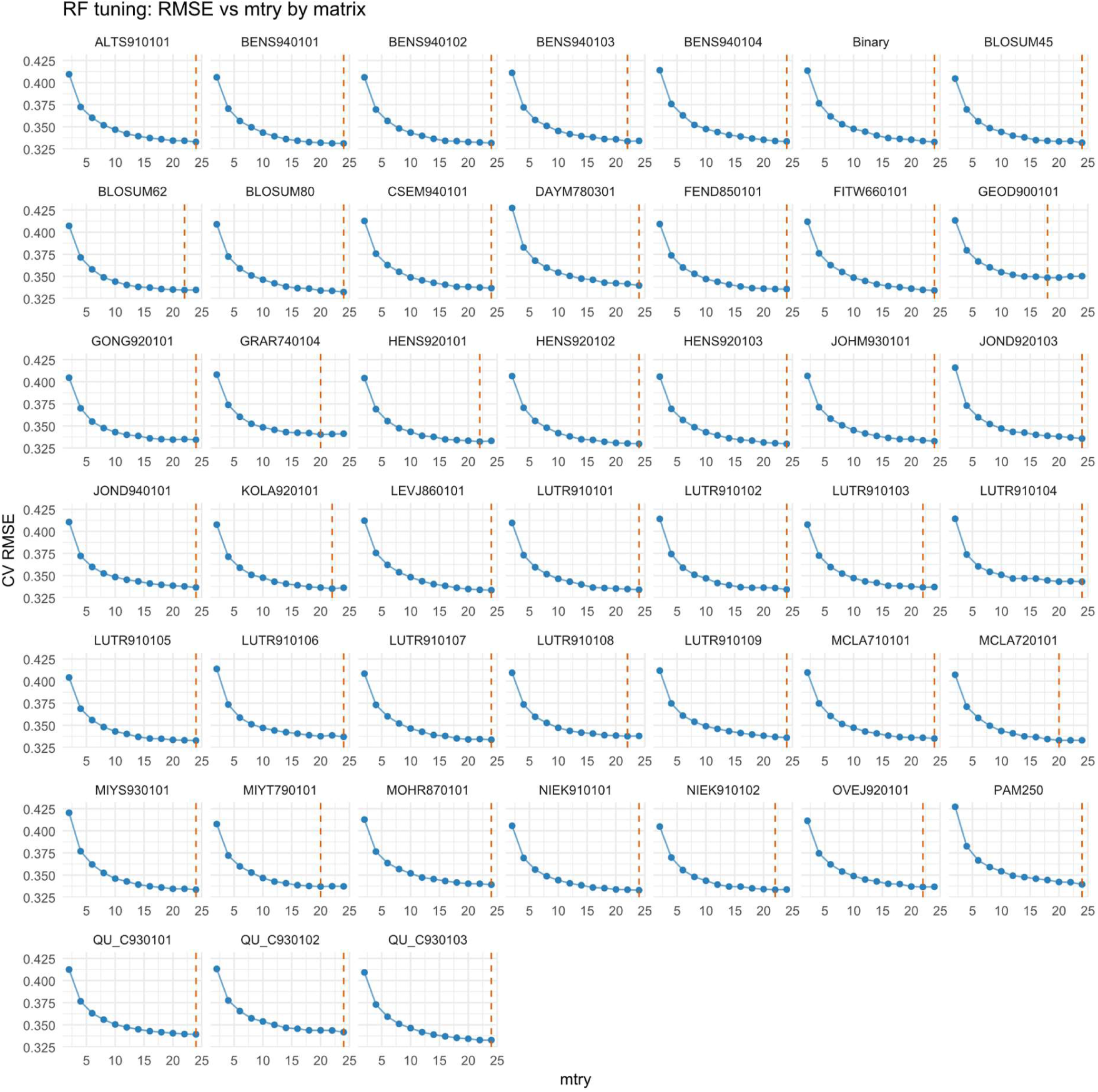
*mtry* optimization. For each substitution matrix, indicated in the figure caption, RMSE was optimized by testing *mtry* values from 1 to 25 and selecting the value that yielded the lowest RMSE.

**Figure 3. Figure supplement 2.**
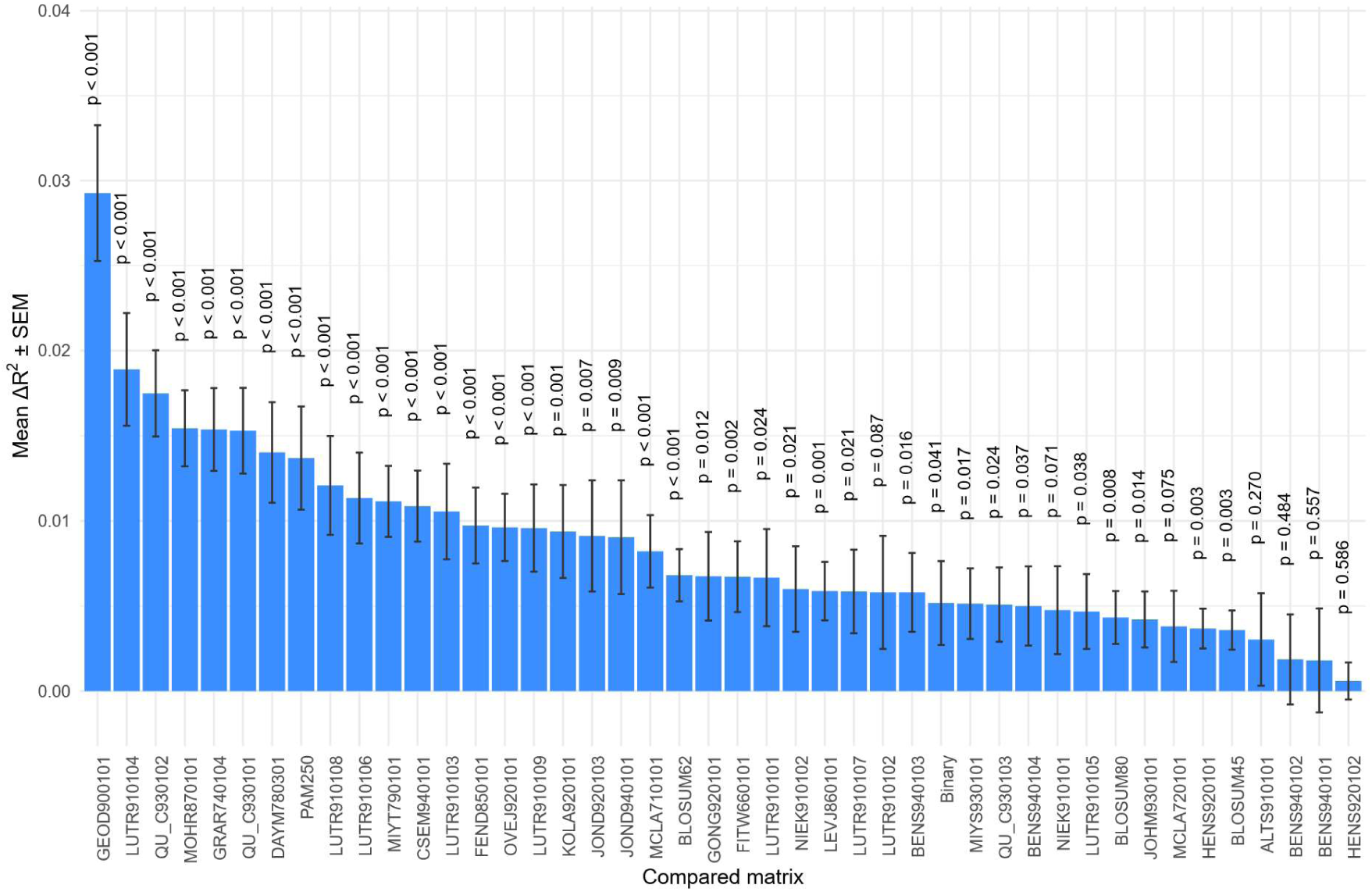
Comparison of the best-performing index matrix with all other matrices. Bars show the delta *R^2^* between predicted and observed antigenic distances across paired cross-validation resamples. Error bars represent the s.e.m. P values were calculated from paired cross-validation comparisons between the top-performing matrix (HENS920103) and each other matrix using a paired t-test.

**Figure 3 – Figure supplement 3.**
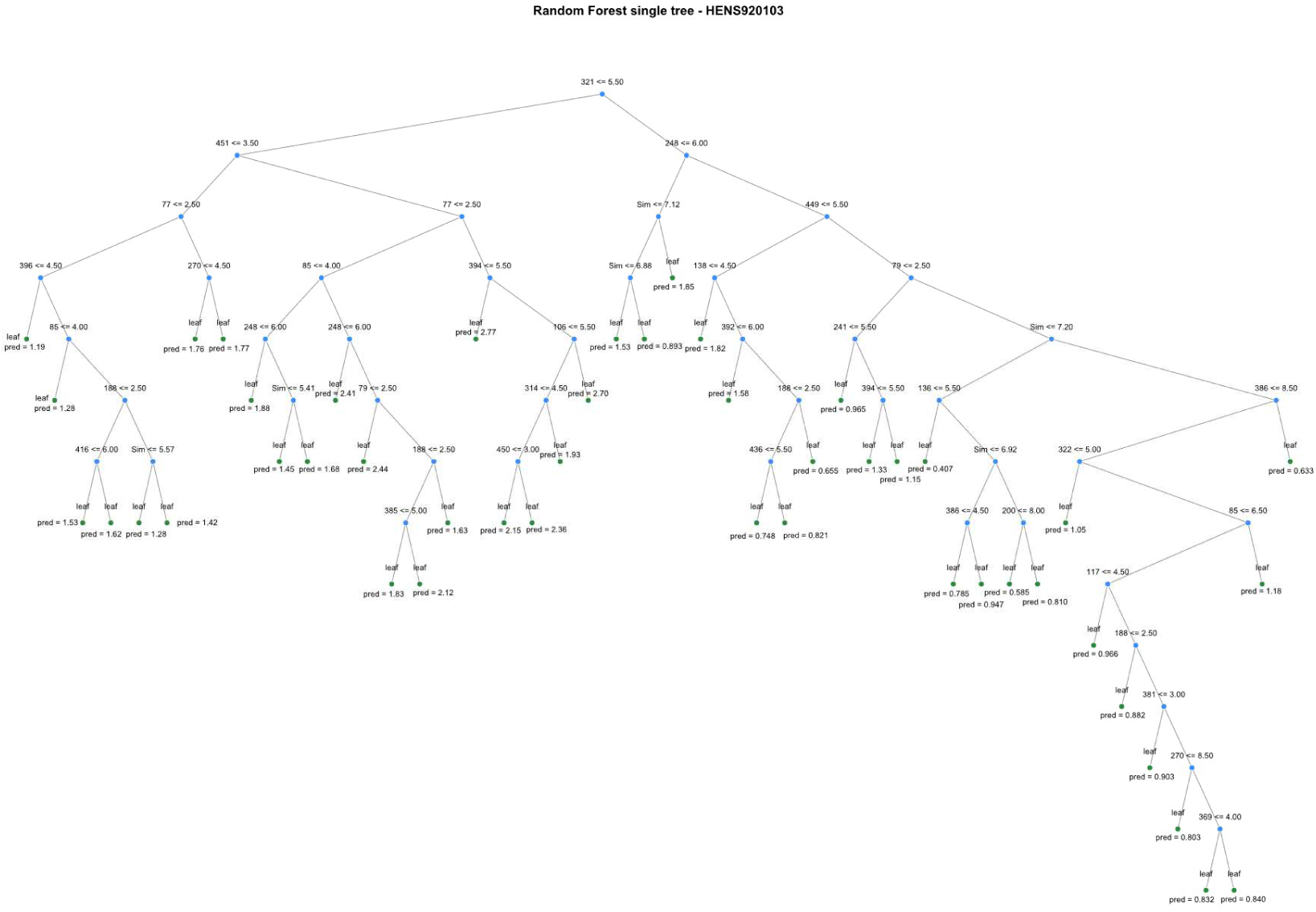
The decision tree of the RF model. The tree represents one of 500 trees generated based on the HENS920103 amino acid index matrix.

**Figure 4 – Figure supplement 1.**
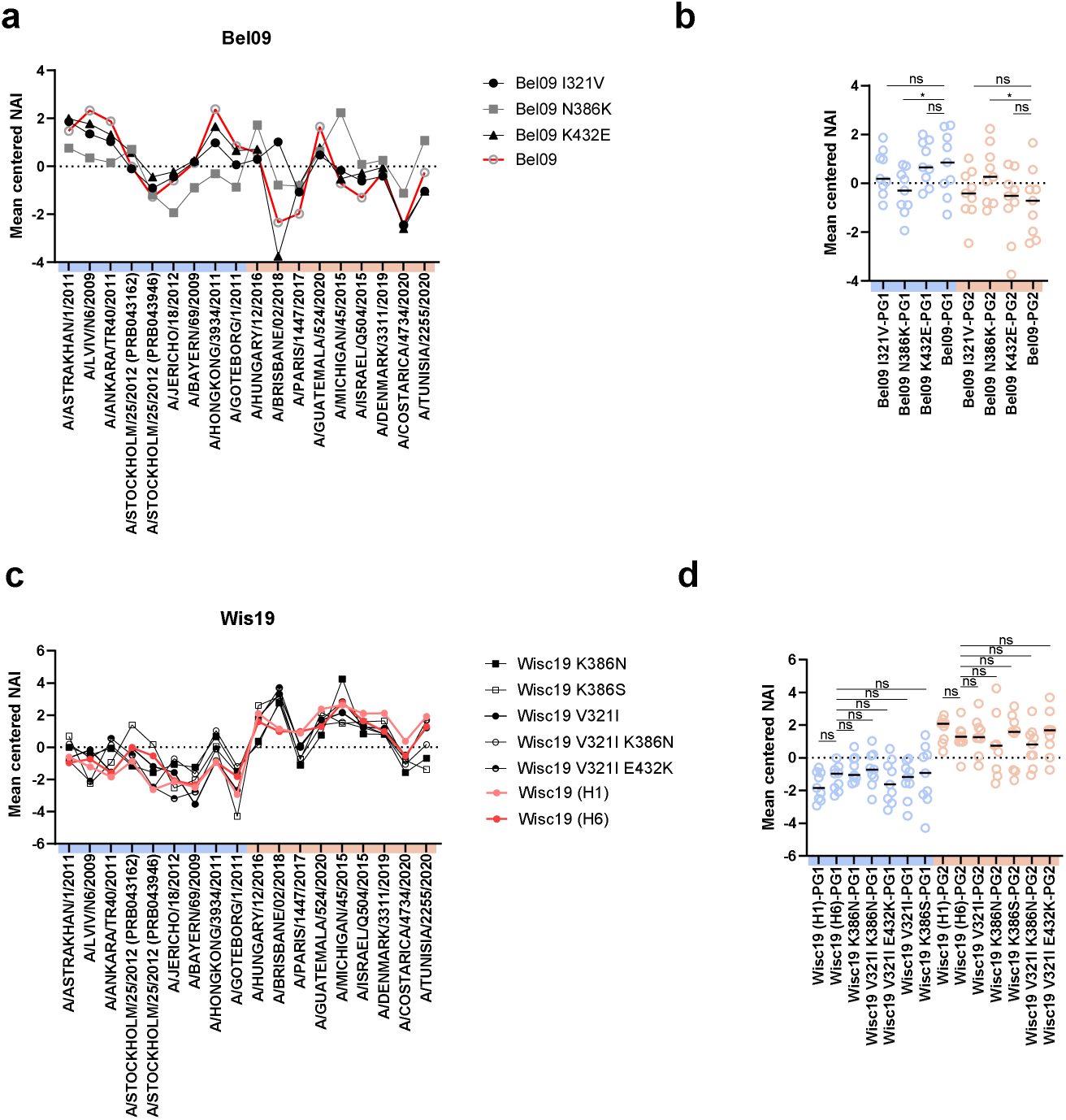
Mean centered NAI determined against mutant H6N1 viruses. (a) H6N_Bel09_ carrying WT NA or NA with the substitutions I321V, N386K, or K432E, tested against the panel of anti-N1 sera. NAI were centered 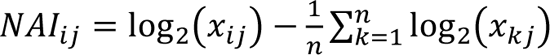. (b) H6N centered means were grouped by phylogenetic group and compared to the parental strain Bel09. (c) HxN_Wis19_ carrying K386N, K386S, V321I, V321I/K386N, V321I/E432K substitutions in N1 or the parental N_Wis19_ (as H6 or H1 references, as shown in Figure 2) have NAI determined against the panel of anti-N1 sera. (d) H6N_Wis19_ centered means were grouped by phylogenetic group and compared to the parental strain Wis19 (H6). Mean centered values were compared using Repeated Measures One-Way Analysis of Variance, followed by Sidak’s post hoc test. *P* values: * *P*<0.05.

**Supplementary table 1.** Amino acid differences in recombinant N1 NAs.

| amino acid at position | 77 | 79 | 81 | 85 | 106 | 117 | 136 | 138 | 170 | 188 | 200 | 241 | 248 | 256 | 264 | 270 | 286 | 288 | 313 |
| --- | --- | --- | --- | --- | --- | --- | --- | --- | --- | --- | --- | --- | --- | --- | --- | --- | --- | --- | --- |
| A/LVIV/N6/2009 | G | S | V | L | I | I | Q | A | Y | Y | N | V | D | F | V | N | S | I | Q |
| A/ASTRAKHAN/1/2011 | . | . | . | . | . | . | . | . | . | . | . | I | . | . | . | . | . | . | . |
| A/ANKARA/TR40/2011 | . | . | A | . | . | . | . | . | H | . | . | I | . | . | . | . | . | . | . |
| A/STOCKHOLM/25/2012 | . | . | . | . | . | . | . | . | . | . | . | I | . | . | . | . | . | . | . |
| A/BAYERN/69/2009 | . | . | . | . | V | . | . | . | . | . | . | . | N | . | . | . | G | . | . |
| A/JERICO/18/2012 | . | . | . | . | V | . | . | . | . | . | S | I | . | L | . | . | . | . | . |
| A/HONGKONG/3934/2011 | . | . | . | . | . | . | . | . | . | . | . | . | . | . | . | . | . | . | R |
| A/GOTEBORG/1/2011 | . | P | . | . | . | . | . | . | . | . | . | . | . | . | . | . | . | . | R |
| A/PARIS/1447/2017 | R | . | A | . | V | . | . | . | . | . | S | I | . | . | I | K | . | . | . |
| A/BRISBANE/02/2018 | R | . | A | . | V | . | . | . | . | T | S | I | . | . | I | K | . | . | . |
| A/GUATEMALA/524/2020 | R | . | A | . | V | . | . | . | . | . | S | I | . | . | I | K | . | . | . |
| A/MICHIGAN/45/2015 | . | . | . | . | V | . | . | . | . | . | S | I | . | . | I | K | . | . | . |
| A/DENMARK/3311/2019 | R | . | A | I | V | . | K | . | . | T | S | I | . | . | I | K | . | . | . |
| A/COSTARICA/4734/2020 | R | . | A | I | V | . | . | S | . | T | S | I | . | . | I | K | . | . | . |
| A/HUNGARY/12/2016 | . | . | . | . | V | M | . | . | . | . | S | I | . | . | . | . | . | V | . |
| A/ISRAEL/Q504/2015 | . | . | . | . | V | . | . | . | . | . | S | I | . | . | I | K | . | . | . |
| A/TUNISIA/2255/2020 | R | . | A | . | V | . | . | . | . | T | S | I | . | . | I | K | . | . | . |

Supplementary table 1 (continued) – Amino acid differences in recombinant N1 NAs.
| amino acid at position | 314 | 321 | 322 | 369 | 381 | 385 | 386 | 389 | 392 | 394 | 396 | 416 | 432 | 436 | 449 | 450 | 451 | 452 | 467 |
| --- | --- | --- | --- | --- | --- | --- | --- | --- | --- | --- | --- | --- | --- | --- | --- | --- | --- | --- | --- |
| A/LVIV/N6/2009 | I | I | F | N | T | S | N | I | D | V | I | D | K | I | N | S | D | T | I |
| A/ASTRAKHAN/1/2011 | . | . | . | K | . | . | . | . | . | . | . | . | . | . | . | . | . | . | . |
| A/ANKARA/TR40/2011 | . | . | . | K | . | . | . | . | . | . | . | . | . | . | . | . | . | . | . |
| A/STOCKHOLM/25/2012 | . | . | . | K | . | . | S | . | . | . | . | . | . | . | . | . | G | . | . |
| A/BAYERN/69/2009 | . | . | . | . | . | . | . | . | . | . | . | . | . | . | . | . | . | . | . |
| A/JERICO/18/2012 | . | . | . | K | . | . | . | . | . | . | . | . | . | . | . | . | . | . | . |
| A/HONGKONG/3934/2011 | . | . | . | . | . | . | . | . | . | I | . | . | . | . | . | . | . | . | V |
| A/GOTEBORG/1/2011 | . | . | . | . | . | . | . | . | . | I | . | . | . | . | . | . | . | . | V |
| A/PARIS/1447/2017 | M | V | . | K | . | . | K | . | E | . | . | . | E | . | D | . | . | . | . |
| A/BRISBANE/02/2018 | M | V | . | K | . | . | K | . | . | . | . | . | E | . | D | . | . | . | . |
| A/GUATEMALA/524/2020 | . | V | L | K | . | . | K | . | . | . | . | . | E | . | D | . | . | . | . |
| A/MICHIGAN/45/2015 | M | V | . | K | . | . | K | . | . | . | . | . | E | . | . | . | . | . | . |
| A/DENMARK/3311/2019 | M | V | . | K | . | . | K | . | . | . | . | . | E | V | D | . | . | . | . |
| A/COSTARICA/4734/2020 | M | V | . | K | . | . | K | . | . | . | M | . | E | V | D | . | . | . | . |
| A/HUNGARY/12/2016 | M | V | . | K | . | . | K | . | . | . | . | . | E | . | . | G | . | . | . |
| A/ISRAEL/Q504/2015 | . | V | . | K | I | . | K | . | . | . | . | . | E | . | . | . | . | . | . |
| A/TUNISIA/2255/2020 | M | V | . | K | . | S | K | K | . | . | . | N | E | . | D | . | . | I | . |

